# MDM2 inhibitors, nutlin-3a and navtemadelin, retain efficacy in human and mouse cancer cells cultured in hypoxia

**DOI:** 10.1101/2022.10.20.513039

**Authors:** Ada Lerma Clavero, Paula Lafqvist Boqvist, Katrine Ingelshed, Cecilia Bosdotter, Saikiran Sedimbi, Long Jiang, Fredrik Wermeling, Borivoj Vojtesek, David P. Lane, Pavitra Kannan

**Affiliations:** Department of Microbiology, Tumor and Cell Biology, Karolinska Institutet, 171 77 Stockholm, Sweden; Department of Medicine Solna, Center for Molecular Medicine, Karolinska University Hospital and Karolinska Institutet, 171 77 Stockholm, Sweden; RECAMO, Masaryk Memorial Cancer Institute, 656 53 Brno, Czech Republic

**Keywords:** p53 activation, MDM2 inhibitors, tumor hypoxia, cancer resistance

## Abstract

Activation of p53 by small molecule MDM2 inhibitors can induce cell cycle arrest or death in p53 wildtype cancer cells. However, cancer cells exposed to hypoxia can develop resistance to other small molecules, such as chemotherapies, that activate p53. Here, we evaluated whether hypoxia could render cancer cells insensitive to two MDM2 inhibitors with different potencies, nutlin-3a and navtemadlin. Inhibitor efficacy and potency were evaluated under short-term hypoxic conditions in human and mouse cancer cells expressing different p53 genotypes (wild-type, mutant, or null). Treatment of wild-type p53 cancer cells with MDM2 inhibitors reduced cell growth by >75% in hypoxia through activation of the p53-p21 signaling pathway; no inhibitor-induced growth reduction was observed in hypoxic mutant or null p53 cells except at very high concentrations. The concentration of inhibitors needed to induce the maximal p53 response was not significantly different in hypoxia compared to normoxia. However, inhibitor efficacy varied by species and by cell line, with stronger effects at lower concentrations observed in human cell lines than in mouse cell lines grown as 2D and 3D cultures. Together, these results indicate that MDM2 inhibitors retain efficacy in hypoxia, suggesting they could be useful for targeting acutely hypoxic cancer cells.

## 1. Introduction

The tumor suppressor protein p53 regulates the cellular response to different types of stress. Under normal (stress-free) conditions, p53 is expressed at a basal level, due to its ubiquitination and proteasomal degradation mediated by the E3 ligases MDM2 and MDMX. However, under stressed conditions, p53 is stabilized and functionally activated, leading to the transactivation of target genes involved in DNA repair, senescence, cell cycle, and/or apoptosis [1]. Given that this response protects cells from damage that can lead to tumor formation, it is not surprising that the *TP53* gene is inactivated or mutated in roughly 50% of all cancers [2]. In the remaining cancers with wildtype p53, protein expression can be silenced through upstream alterations. Restoration of wildtype p53 activity in tumors is therefore an important goal for improved cancer treatment [3].

Several small molecules have been developed to restore p53 activity in cancer cells expressing wildtype p53. One class of these molecules, the MDM2 inhibitors, prevent the interaction between p53 and its negative regulator MDM2. As a result, p53 accumulates in the cell and activates its downstream target genes [4]. While treatment with inhibitors alone tends to induce cell cycle arrest, treatment with inhibitors combined with chemotherapy or radiation tends to enhance apoptosis *in vitro* and in preclinical tumor models [5–10], although with varying potencies. For example, one of the most studied MDM2 inhibitors, nutlin-3a, requires 2 to 3-fold higher concentrations to induce p53 than other inhibitors (e.g., navtemadlin) require [6,11,12], and may have off-target effects [12]. Although many of these inhibitors show great promise in preclinical and clinical trials, the development of resistance is a concern [13].

Hypoxia, which commonly occurs in solid tumors [14], can render cancer cells resistant to chemotherapies that induce their apoptotic activity through p53-dependent mechanisms [15,16]. This resistance is thought to arise partly from transcriptional and/or translational changes that cells undergo to survive hypoxia [17]. For example, upregulation of hypoxia responsive miRNAs has been shown to render colorectal cancer cells resistant to treatment with 5-fluorouracil [18]. Even 4 h exposure to hypoxia has been shown to reduce ribosome translation in cancer cells [19], which could in turn affect p53 transcriptional activity [20]. In addition, hypoxia may select for cells that have reduced potential for p53-mediated apoptosis [21], potentially affecting the ability of small molecules to activate a p53-mediated response. Moreover, hypoxia has been reported to reduce cell proliferation [22], which could spare hypoxic cells from the cytotoxic effects of chemotherapies targeting proliferating cells. Since hypoxia-mediated resistance is associated with poorer clinical prognosis in several solid tumors [23], evaluating the ability of MDM2 inhibitors to induce a p53 response in hypoxia is important for their successful clinical use.

Although many such inhibitors have been developed, few have been evaluated for their efficacy in hypoxia. One study found that the small molecule RITA, which binds directly to the N-terminus of p53, induced p53-dependent apoptosis in human colorectal cancer cell lines cultured in hypoxia [24]. However, RITA has also been reported to exert apoptosis in a non-p53 dependent manner [25]. Another study reported that the MDM2 inhibitor nutlin-3 activated p53 and inhibited growth in mouse melanoma cells cultured in hypoxia [26]. However, nutlin-3 and its more active enantiomer nutlin-3a have poor bioavailability for clinical use. To our knowledge, no studies have evaluated the efficacy of more potent and bioavailable MDM2 inhibitors, such as navtemadlin (previously known as AMG 232), in hypoxic cancer cells.

Here, we evaluated the ability of two MDM2 inhibitors with different potencies, nutlin-3a and navtemadlin, to induce p53 levels in human and mouse tumor cells cultured in hypoxia. Changes in cell proliferation were quantified using growth assays and flow cytometry, while molecular changes in the p53-p21 axis were detected using transcriptional and translational assays. Our results show that both inhibitors induce a p53 response in hypoxic cancer cells with wildtype p53, but with varying efficacies depending on species and cell line.

## 2. Materials and methods

### 2.1. Chemicals

Nutlin-3a (Sigma Aldrich), navtemadlin (2639, Axon Medchem), and staurosporine (Sigma Aldrich) were dissolved in DMSO (D2650, Sigma Aldrich). EF5 (Merck Millipore) was dissolved in 5% glucose in sterile saline, as per manufacturer’s instructions.

### 2.2. Cell culture

Six cell lines were used: HCT116 p53^+/+^ and HCT116 p53^−/−^ (human colorectal), MCF7 (human breast adenocarcinoma), B16-F10 p53^+/+^ and B16-F10 p53^−/−^ (mouse melanoma), and HT29 (human colorectal with mutant p53). The HCT116 cell lines were kindly provided by Professor Bert Vogelstein (Johns Hopkins University). The B16-F10 p53^+/+^ cell line (ATCC) was used to generate the p53^−/−^ cell line using CRISPR/Cas9 as previously described [27]. MCF7 and HT29 were obtained from ATCC and used within 6 months of purchase. HCT116 and HT29 cells were cultured in 1.0 g/L glucose DMEM (31885049, ThermoFisher), MCF7 cells in 1.0 g/L glucose MEM (31095029, ThermoFisher), and B16-F10 cells in no glucose RPMI (11879020, ThermoFisher) supplemented with 1.0 g/L glucose (A2494001, ThermoFisher). All media were supplemented with 10% heat inactivated fetal bovine serum (RC35964, HyClone), 1% penicillin/streptomycin (P0781, Sigma-Aldrich), and 10 mM HEPES (H3537, Sigma-Aldrich). Culture medium was changed every 2-3 days. Cells were maintained in an incubator at 37 ^∘^C, 5% CO_2_, and 21% O_2_, or in an Invivo2 chamber (Baker Ruskinn 400) for hypoxia at 37 ^∘^C, 5% CO_2_, and 1% O_2_. All cell lines were regularly tested for mycoplasma (Mycoplasma Alert, Lonza).

### 2.3. Cellular growth inhibition

The ability of MDM2 inhibitors to inhibit cell growth in normoxia and hypoxia was measured using a modified version of a cellular viability assay [28]. Single cells (5×10^3^ cells in 100 *μ*L/well) were seeded in 96-well plates and allowed to attach for 24 h. Stock solutions of nutlin-3a and navtemadlin were prepared in growth medium at 2X the final concentration (50 *μ*M for nutlin-3a; 20 *μ*M for navtemadlin), and serially diluted 1:1 in fresh medium using a 12-well dilution reservoir (VWR) to achieve 10 drug concentrations (including one untreated). Drug-containing medium (100 *μ*L) was then added to each well to achieve the final 1X concentration. After cells were incubated for 72 hours in normoxia or hypoxia, they were fixed with 10% neutral buffered formalin (Sigma) for 15 min and imaged in PBS (100 *μ*L/well) using brightfield microscopy (Incucyte^®^ S3 Live-Cell Analysis System, Essen Bioscience). Cell confluence was determined using the Incucyte software (Segmentation value: 0.4 for HCT116 and B16-F10 cells and 1.2 for MCF7 cells; Hole fill: 200 μm^3^; Minimum area: 200 μm^3^). The concentration of drug that reduced cell growth by 50% of the untreated control (IC_50_) was calculated using GraphPad Prism (Nonlinear fit equation: inhibitor vs response—variable slope).

### 2.4. Cell cycle analysis

The effect of MDM2 inhibitors on cell cycle distribution was assessed using flow cytometry. Cells (1×10^5^) were seeded in T-25 culture flasks and incubated for 72 hours prior to treatment. After the medium in each flask was replaced with fresh medium (5 mL/flask) containing <0.1% DMSO, 2 *μ*M nutlin-3a, or 0.5 *μ*M navtemadlin, cells were incubated either in normoxia or in hypoxia for a total of 24h. One hour prior to trypsin-mediated harvest, cells were treated with 10 *μ*M of EdU (5-ethynyl-2′-deoxyuridine) to mark cells in S phase. Cells were then fixed by adding 100 *μ*L of Click-iT^®^ fixative (Click-iT™ Plus EdU Alexa Fluor™ 647 Flow Cytometry Assay Kit, ThermoFisher) and stored at 4°C for 24 hours. Following fixation, cells were permeabilized using Click-iT^®^ saponin-based permeabilization and wash buffer, and stained with Alexa Fluor^®^ 647 picolyl azide, according to manufacturer’s instructions. Cells were suspended in 1X saponin buffer containing FxCycle Violet (1:1000, Thermo Scientific) to stain DNA. Cell cycle distribution was measured using the FACS Canto II or BD LSR II flow cytometers. Data were analyzed using FlowJo software (v. 10).

### 2.5. qPCR

Gene expression induced by MDM2 inhibitors in normoxia and hypoxia was measured using qPCR. Cells (1×10^5^ cells in 2 mL/well) were seeded in a 6-well tissue culture dish (TPP) and allowed to attach for 36 h. Drug treatment was carried out by replacing the medium with fresh medium containing <0.1% DMSO, 2 *μ*M nutlin-3a, or 0.5 *μ*M navtemadlin, and incubating cells in normoxia or hypoxia for 24 h. Cells were lysed using RNA binding lysis buffer (500 *μ*L/well; mirVana™ miRNA Isolation Kit, ThermoFisher), and lysates were stored at −80 ^∘^C until extraction. Total RNA was extracted using the manufacturer’s protocol (mirVana™ miRNA Isolation Kit, ThermoFisher Scientific or RNAeasy, Qiagen) and stored at −20 ^∘^C. cDNA was obtained by reverse transcription (iScript cDNA Synthesis Kit, BIO RAD). qPCR was performed in duplicate in a 96-well plate according to manufacturer’s instructions (iTaq Universal SYBR Green Supermix, BioRad) using pre-designed, PrimeTime qPCR primers containing SYBR Green (Integrated DNA Technologies; Table1). Primers were confirmed to have 90-110% efficiency using the standard curve method. Gene expression was measured for two target genes (*CDKN1A* and *MDM2* for human; *Cdkn1a* and *Mdm2* for mouse) and reference genes (*B2M* for human and *Rplp0* for mouse). StepOnePlus™ Software v2.3 was used to analyze the data and fold change was calculated using the 2-⊿⊿Ct method, with values for target genes normalized to the reference genes.

**Table 1.**
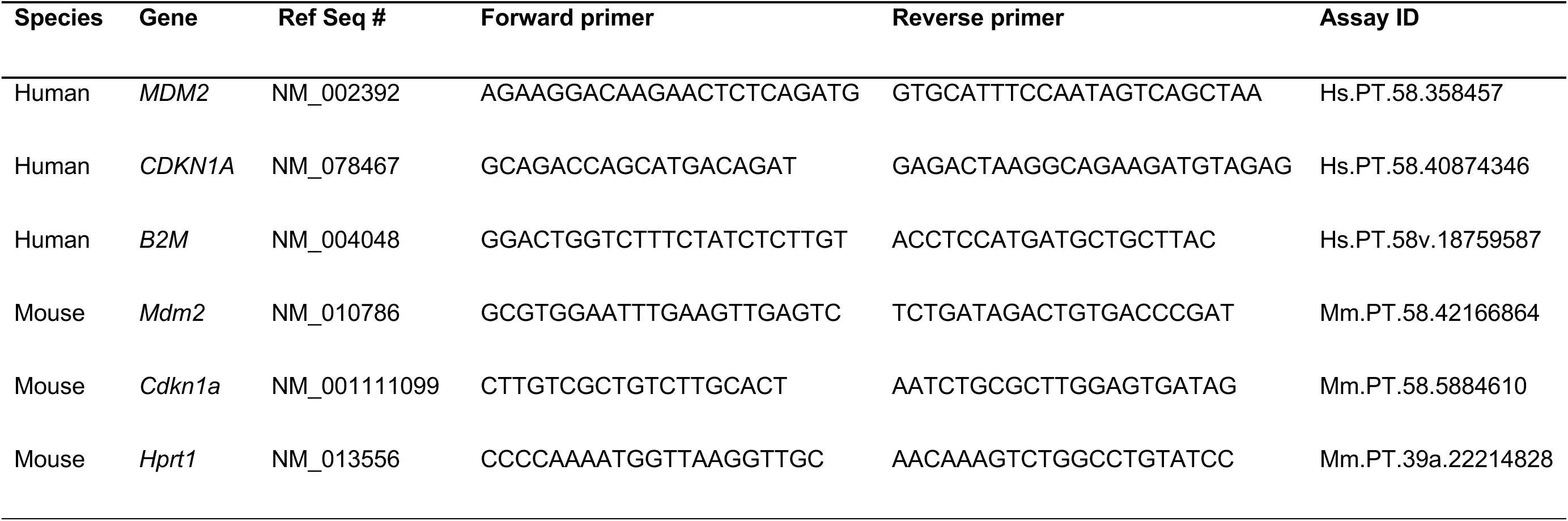
Primer sequences used for quantitative PCR to measure expression of genes in human and mouse cancer cell lines.

### 2.6. Western blotting

To determine whether MDM2 inhibitors induced known changes in p53-p21 axis at the protein level in hypoxia, we examined p53, p21, MDM2 and HIF1a levels in normoxia and hypoxia using immunoblotting. Cells (4×10^5^ cells in 3 mL/dish) were seeded in a 60 mm tissue culture dish (TPP) and allowed to attach for 36 h. Drug treatment was carried out by replacing the medium with fresh medium containing <0.1% DMSO, 2 *μ*M nutlin-3a, or 0.5 *μ*M navtemadlin, and incubating cells in normoxia or hypoxia for 24 h. Adherent cells were washed with PBS, lysed with SDS lysis buffer (1X SDS lysis buffer, BioRad), heated at 95 ^∘^C for 5 min, sonicated (Qsonica Sonicators) for 30 sec at 20% amplitude, and stored at −20 ^∘^C until use.

Prior to electrophoresis, protein concentrations were determined using the DC Protein Assay (BioRad) according to manufacturer’s instructions. Equal amounts of protein (15 *μ*g) were loaded on 4-15% 1 D polyacrylamide Mini-PROTEAN TGX Stain-Free gels (BioRad) and run in 1X Tris Glycine SDS buffer (BioRad) at 50 V for 5 min and 150 V for 45 min. Gels were activated by a 5 min exposure using an Imaging System (ChemiDoc^TM^ Touch, BioRad). Proteins were then transferred to a Minisize PVDF 7 membrane (BioRad) using a semi-dry transfer (TurboBlot, BioRad) for 30 min using the standard built-in protocol. Membranes were imaged using the Imaging System (ChemiDoc^TM^ Touch, BioRad) to obtain images of total protein and subsequently equilibrated in PBS-T (0.1% Tween-20 in PBS) for 15 min. Membranes were subsequently incubated in PBS-T containing 5% milk for 1 h at room temperature to block non-specific binding. Membranes were then incubated with primary antibodies overnight at 4 ^∘^C (Table 2). The next day, membranes were washed 4 times for 5 min in PBS-T and then incubated for 1 h room temperature with secondary antibodies (Table 2). After membranes were washed 6 times for 5 min in PBS-T, they were incubated with ECL Clarity Substrate and Reagent (Clarity™ Western ECL Substrate, 1705060, BioRad) for 5 min and imaged for chemiluminescence signal (ChemiDoc^TM^ Touch, BioRad).

**Table 2.** Dilutions of 1 and 2 antibodies used for immunoblotting detection of proteins involved in hypoxia and p53 activation in human and mouse cancer cell lysates. HIF1α = hypoxia inducible factor 1α; Hs = human; m =mouse

| Protein target | Primary antibody |  |  |  | Secondary antibody |  |  |  |
| --- | --- | --- | --- | --- | --- | --- | --- | --- |
|  | 1° Antibody (clone or catalog) | Host species | Company | Dilution of 1° | 2° Antibody | Host species | Company | Dilution of 2° |
| HIF1α (Hs) | 54/MOP1 | Mouse | BD biosciences | 1: 500 | Anti-mouse HRP | Goat | Dako | 1: 1000 |
| p53 (Hs) | DO-1 | Mouse | In house[48] | 1: 3000 | Anti-mouse HRP | Goat | Dako | 1: 2000 |
| p21 (Hs) | SX-118 | Mouse | Santa Cruz | 1: 500 | Anti-mouse HRP | Goat | Dako | 1: 1000 |
| HIF1α (m) | NB100-449 | Rabbit | BD biosciences | 1: 200 | Anti-rabbit HRP | Swine | Dako | 1: 1000 |
| p53 (m) | EPR20416-124 | Rabbit | Abcam | 1: 1000 | Anti-rabbit HRP | Swine | Dako | 1: 1000 |
| p21 (m) | EPR18021 | Rabbit | Abcam | 1: 1000 | Anti-rabbit HRP | Swine | Dako | 1: 1000 |

### 2.7. Spheroid growth assays

To determine whether MDM2 inhibitors could reduce growth in 3D tumors comprising innate hypoxia, we measured the growth of 3D spheroids under treatment. Single cell solutions (3×10^3^ cells in 100 μL/well) were seeded in ultra-low attachment, round-bottom, 96-well plates to generate spheroids (Corning 7007). After the formation of spheroids (72 h), fresh medium (100 μL) containing twice the final concentration of drug was added to each well resulting in a total volume of 200 μL. Spheroids from human cells were treated with a final concentration of 5 μM nutlin-3a, 1 μM navtemadlin, and 2 μM staurosporine, while those from mouse cells were treated with a final concentration of 10 μM nutlin-3a, 2 μM navtemadlin, and 5 μM staurosporine. Half the medium was replaced with fresh medium every two days. Spheroids were imaged for up to 4-7 days (Biospa 8 or Incucyte Software). Images were analyzed using SpheroidSizer [29].

### 2.8. Statistical analysis

After checking for homogeneity of variance, data were evaluated for statistical significance using a mixed-effect, two-way ANOVA with Dunnett’s multiple comparison’s test (unpaired, two-tailed, *α* = 0.05). To avoid violating assumptions in normality, statistical significance for qPCR data was calculated on dCt values (the difference between sample and reference Ct values). For monolayer assays, data points represent the average value from ≥3 biological experiments. For spheroid assays, data points represent individual spheroids combined from ≥2 biological experiments. Randomization and blinding were not possible.

## 3. Results

### 3.1. Nutlin-3a and navtemadlin reduce growth of p53^WT^ cancer cells in hypoxia

In normoxia (21% O_2_) and hypoxia (1% O_2_), treatment with the MDM2 inhibitors, nutlin-3a and navtemadlin, reduced cell growth in three p53 wildtype (p53^WT^) cell lines (human colorectal carcinoma HCT116 p53^+/+^; human breast carcinoma MCF7; mouse melanoma B16-F10 p53^+/+^) (Fig. 1a-c). In normoxia, both inhibitors reduced cell growth by a maximum of 75%, with IC50 values ranging from 1.6 – 8.6 μM for nutlin-3a and 0.2 – 1.4 μM for navtemadlin. Similar reductions in cell growth were observed in hypoxia, with IC50 values ranging from 1.4 – 6.7 μM for nutlin-3a and 0.3 – 1.3 μM for navtemadlin (Table 3). In contrast, except at very high concentrations of the drugs, no reductions in cell growth were measured in p53 knockout (p53^KO^; HCT116 p53^−/−^; B16-F10 p53^−/−^) and p53 mutant (p53^mut^; human colorectal carcinoma HT-29) cell lines treated in normoxia or hypoxia (Fig 1d-f), confirming the wildtype p53-dependent mechanism of action of the inhibitors. Together, these findings indicate that the potencies of nutlin-3a and navtemadlin are not significantly affected in hypoxic conditions, and they confirm that nutlin-3a is less potent than navtemadlin in normoxia and hypoxia.

**Table 3.** Potencies of nutlin-3a and navtemadlin measured under normoxic and hypoxic conditions in cancer cell lines expressing wild-type p53. Potency was calculated as the concentration of drug that reduced cell growth by 50% of the untreated control (IC_50_), as measured by a cellular viability assay. Cellular confluence was measured after 72 h of exposure to 10 drug concentrations. CI represents the confidence interval.

| Cell line | nutlin-3a |  |  | navtemadlin |  |  |
| --- | --- | --- | --- | --- | --- | --- |
|  | Normoxia IC<sub>50</sub> | Hypoxia IC<sub>50</sub> | *P*<sub>adj</sub> | Normoxia IC<sub>50</sub> | Hypoxia IC<sub>50</sub> | *P*<sub>adj</sub> |
|  | (95% CI) | (95% CI) |  | (95% CI) | (95% CI) |  |
| HCT116 p53<sup>+/+</sup> | 1.6 (1.3 to 1.9) | 1.4 (1.1 to 1.6) | 0.99 | 0.3 (0.3 to 0.4) | 0.3 (0.2 to 0.4) | 0.99 |
| MCF7 | 1.8 (0.9 to 2.8) | 2.7 (0 to 8.8) | 0.96 | 0.2 (0.2 to 0.3) | 0.3 (0.1 to 0.4) | 0.95 |
| B16-F10 p53<sup>+/+</sup> | 8.6 (6.9 to 10.3) | 6.9 (3.2 to 10.2) | 0.74 | 1.4 (1.2 to 1.5) | 1.3 (1.0 to 1.7) | 0.96 |

**Figure 1.**
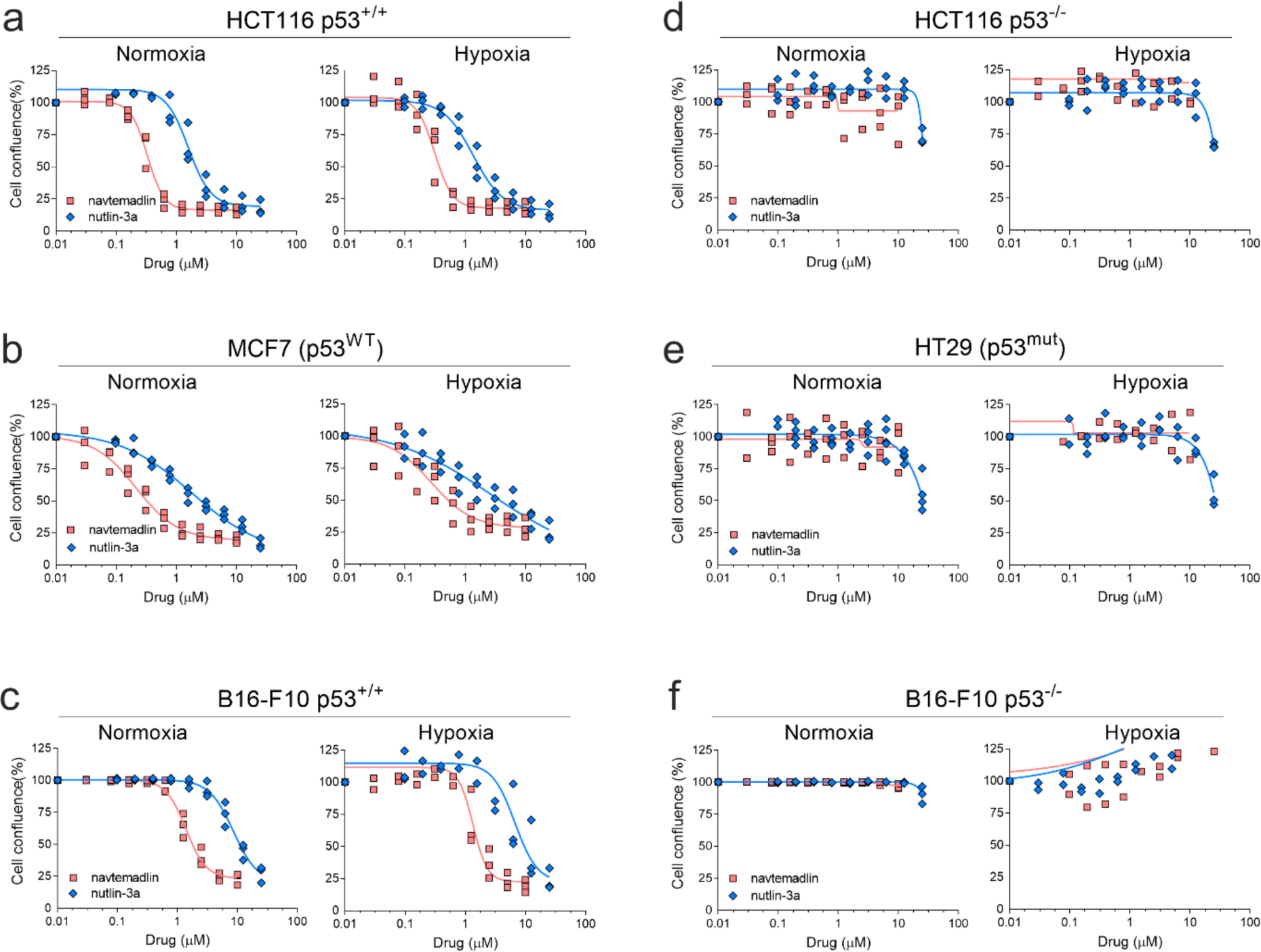
MDM2 inhibitors, nutlin-3a and navtemadlin, inhibit the growth of p53 wildtype (p53^wt^) cancer cell lines cultured in hypoxia (1% O_2_). (**a-c)** Concentration-growth curves of three p53 wildtype cell lines (human HCT116 p53^+/+^, human MCF7, and mouse B16-F10 p53^+/+^) in normoxic and hypoxic conditions. **(d-f)** Concentration-growth curves of three p53 knockout and mutant cell lines (human HCT116 p53^−/−^, human HT-29 p53^mut^, and mouse B16-F10 p53^−/−^) in normoxic and hypoxic conditions. Cell growth (as assessed by percent confluence) was measured after 72 h of treatment with MDM2 inhibitors. Each data point represents the averaged value of triplicates from one independent experiment (n = three independent experiments). The line indicates the best fit model for concentration-growth response.

To determine whether the reduction in growth resulted from fewer proliferating cells, we then measured the uptake of 5-ethynyl-2’-deoxyuridine (a marker of DNA synthesis) in cells treated with MDM2 inhibitors for 24 h in normoxia or hypoxia. In HCT116 p53^+/+^ cells, treatment with both inhibitors in normoxia led to a > 60% decrease in S phase (nutlin-3a, *P_adj_* = 0.0016; navtemadlin, *P_adj_* < 0.0001; Fig 2a; SI Fig 1). In hypoxia, however, only treatment with navtemadlin led to a significant decrease in S phase (nutlin-3a, *Padj* = 0.08; navtemadlin, *P_adj_* < 0.0001; Fig 2a). Although hypoxia itself is reported to reduce cell proliferation and induce cell cycle arrest [22], it alone did not affect proliferation after 24 h in HCT116 p53^+/+^ cells (S phase, DMSO normoxia vs hypoxia, *P_adj_* = 0.92). No significant changes in cell proliferation were measured for HCT116 p53^−/−^ cell lines treated in either normoxia or hypoxia (Fig 2b). Similarly, in B16-F10 p53^+/+^ cells, but not B16-F10 p53^−/−^ cells, drug treatment in normoxia led to a reduction > 90% decrease in S phase (nutlin-3a, *P_adj_* = 0.0003; navtemadlin, *P_adj_* = 0.0007; Fig 2c). However, hypoxia alone reduced the proliferation of B16-F10 p53^+/+^ by 83% (S phase, DMSO normoxia vs hypoxia, *P_adj_* = 0.003) and that of B16-F10 p53^−/−^ by 75% (S phase, DMSO normoxia vs hypoxia, *P_adj_* = 0.03; Fig 2d). Treatment with either inhibitor in hypoxia did not lead to a further decrease in S phase of B16-F10 p53^+/+^ (nutlin-3a, *P_adj_* = 0.96; navtemadlin, *P_adj_* = 0.97; Fig 2c) or of B16-F10 p53^−/−^ cells (nutlin-3a, *P_adj_* = 0.99; navtemadlin, *P_adj_* = 0.99; Fig 2d). These results suggest that while MDM2 inhibitors reduce proliferation of human and mouse cells in normoxia, they may not further reduce proliferation in hypoxia if hypoxia alone reduces proliferation.

**Figure 2.**
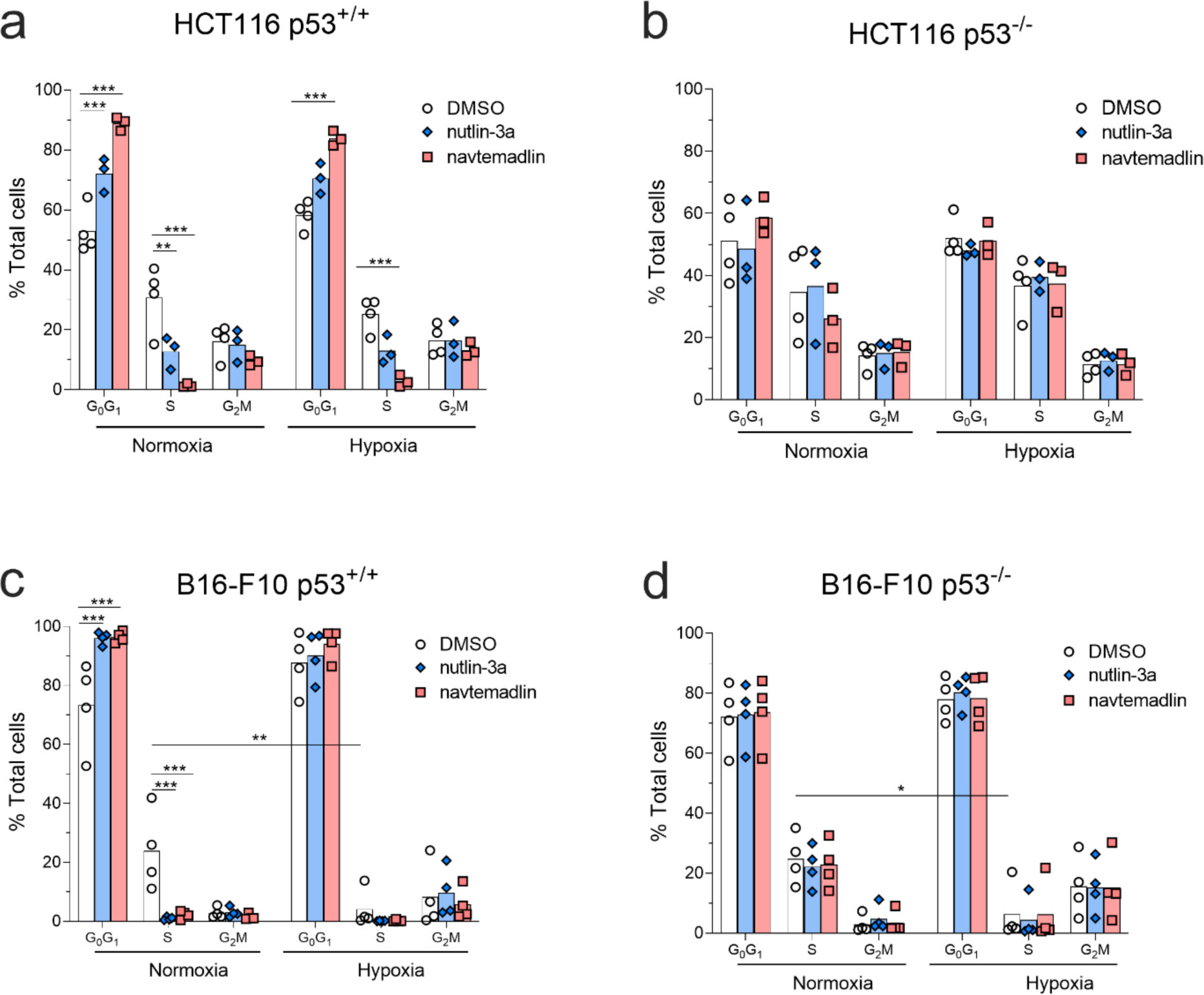
MDM2 inhibitors reduce proliferation of human p53^wt^ cancer cells cultured under hypoxia, while hypoxia alone reduces proliferation of mouse cancer cells. **(a-b)** Cell cycle distribution of human HCT116 p53^+/+^ and p53^−/−^ cells after 24 h treatment with DMSO, nutlin-3a (2 μM), or navtemadlin (0.5 μM) in normoxia or hypoxia. **(c-d)** Cell cycle distribution of mouse B16-F10 p53^+/+^ and p53^−/−^ cells after 24 h treatment with DMSO, nutlin-3a (10 μM), or navtemadlin (2 μM) in normoxia or hypoxia. Bar graphs represent the mean value from four independent experiments; individual data points represent mean value from one experiment. Statistical significance was assessed by 2-way ANOVA followed by post-hoc Sidak’s or Tukey’s correction for multiple testing (unpaired, two-tailed, α = 0.05). Adjusted *P*-values from the post-hoc t-tests are indicated (\**P* < 0.05, \*\**P* < 0.01, \*\*\**P* < 0.001).

### 3.2. Nutlin-3a and navtemadlin activate the p53-p21 axis in hypoxic p53wt cells

Given that MDM2 inhibitors are known to induce cell cycle arrest through activation of the p53 pathway [4], we subsequently measured the mRNA and protein levels of two downstream targets of p53 activated by these inhibitors: MDM2 (negative regulator of p53) and p21 (a marker of cell cycle arrest). In HCT116 p53^+/+^ cells treated with nutlin-3a and navtemadlin for 24 h, the gene expression of *MDM2* and of *CDKN1A* significantly increased by more than 9-fold both in normoxia (*MDM2*: nutlin-3a, *P_adj_* < 0.001; navtemadlin, *P_adj_* < 0.001; *CDKN1A*: nutlin-3a, *P_adj_* < 0.001; navtemadlin, *P_adj_* < 0.001) and in hypoxia (*MDM2*: nutlin-3a, *P_adj_* < 0.001; navtemadlin, *P_adj_* < 0.001; *CDKN1A*: nutlin-3a, *P_adj_* < 0.001; navtemadlin, *P_adj_* < 0.001; Fig 3a). The expression of *CDKN1A* was not enhanced in solvent-treated hypoxic HCT116 p53^+/+^ cells (DMSO normoxia vs hypoxia, *P* = 0.56 by unpaired t-test, two-tailed), corroborating the results from flow cytometry that showed that HCT116 p53^+/+^ cells are not arrested by hypoxia alone. No significant changes in gene expression were measured for HCT116 p53^−/−^ cells in normoxia or hypoxia (*P_adj_* > 0.60 for all comparisons; Fig 3b).

**Figure 3.**
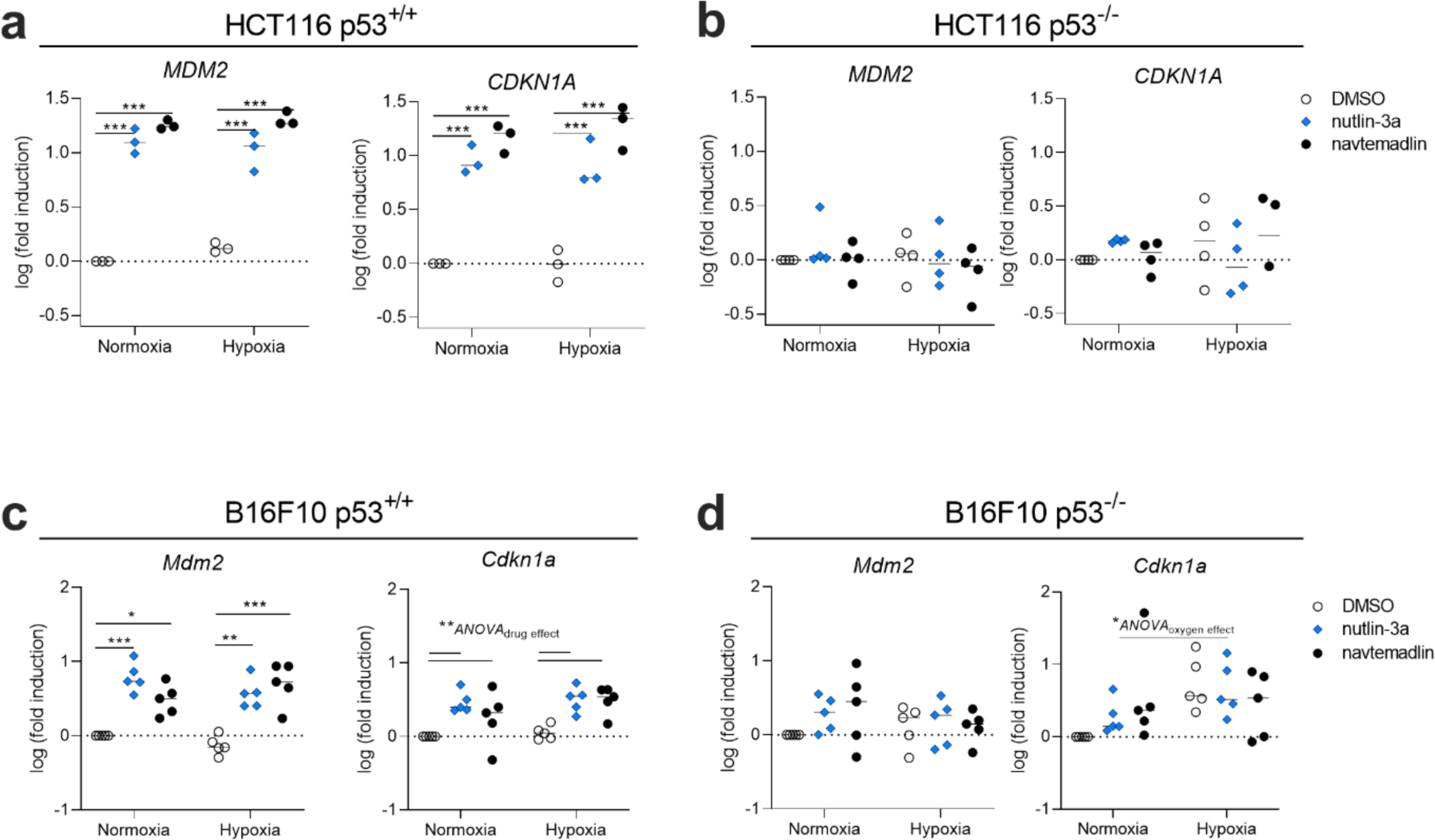
MDM2 inhibitors increase expression of downstream target genes of p53 in hypoxic p53^wt^ cancer cells. Gene expression was measured by qPCR in (**a-b**) HCT116 p53^+/+^ and p53^−/−^ cells, and in (**c-d**) B16-F10 p53^+/+^ and p53^−/−^ cells after 24 h treatment with MDM2 inhibitors. Each data point represents averaged values of duplicates from three to five independent experiments. Expression was normalized to house-keeping genes *B2M* or *Rplp0* for human or mouse cell lines, respectively. Statistical significance was assessed on non-log transformed dCT values using 2-way ANOVA followed by post-hoc Dunnett’s correction for multiple testing (unpaired, two-tailed, α = 0.05). Adjusted *P*-values for individual comparisons (e.g., DMSO vs nutlin-3a) from the Dunnett’s test are indicated (\**P* < 0.05, \*\**P* < 0.01, \*\*\**P* < 0.001). For *Cdkn1a* in **c** and **d**, values indicate the source of variation identified by the ANOVA (run prior to the post-hoc t-test); post-hoc comparisons of each drug against DMSO did not provide significant values, likely due to insufficient power.

In B16-F10 p53^+/+^ cells treated with MDM2 inhibitors for 24 h, the expression of *Mdm2* increased by more than 3-fold in normoxia (nutlin-3a, *P_adj_* < 0.001; navtemadlin, *P_adj_* = 0.02) and in hypoxia (nutlin-3a, *P_adj_* = 0.001; navtemadlin, *P_adj_* < 0.001; Fig 3c). Inhibitor treatment also significantly increased *Cdkn1a* expression (2-way ANOVA interaction model, *Source of variation_(drug)_* = 0.009); however, we likely did not have sufficient power using post-ANOVA t-tests (corrected for multiple testing) to detect significance for the effect of the individual drugs in either normoxia (nutlin-3a, *P_adj_* = 0.05; navtemadlin, *P_adj_* = 0.36) or hypoxia (nutlin-3a, *P_adj_* = 0.06; navtemadlin, *P_adj_* = 0.07; Fig 3c). No significant changes in gene expression were measured for B16-F10 p53^−/−^ cells in normoxia or hypoxia (*P_adj_* > 0.10 for all comparisons; Fig 3d). These results indicate that transcriptional activation by the MDM2 inhibitors occurs with similar efficacy in hypoxia as in normoxia for both human and mouse cell lines.

Protein activation by MDM2 inhibitors was also unaffected by hypoxia. In p53^wt^ cell lines (HCT116 p53^+/+^, MCF7, and B16-F10 p53^+/+^), treatment nutlin-3a and navtemadlin induced p53 and p21 expression in both normoxia and hypoxia (Fig 4), as assessed by immunoblotting. All cell lines cultured in hypoxic conditions expressed the hypoxia induced transcription factor HIF1*α*, confirming that the cells had become hypoxic. Interestingly, HIF1*α* expression decreased upon treatment with nutlin-3a and navtemadlin in MCF7 cell line but did not change in the other cell lines. In HCT116 p53^−/−^ cells, no increase in p53 or p21 expression was detected with drug treatment in normoxia or hypoxia (Fig 4). Similar results were observed in B16-F10 p53^−/−^ cells, even though p21 expression was detected in hypoxia. Together, these data confirm that MDM2 inhibitors activate the p53-p21 axis with similar efficacy in monolayer cultures exposed to hypoxia as to those exposed to normoxia.

**Figure 4.**
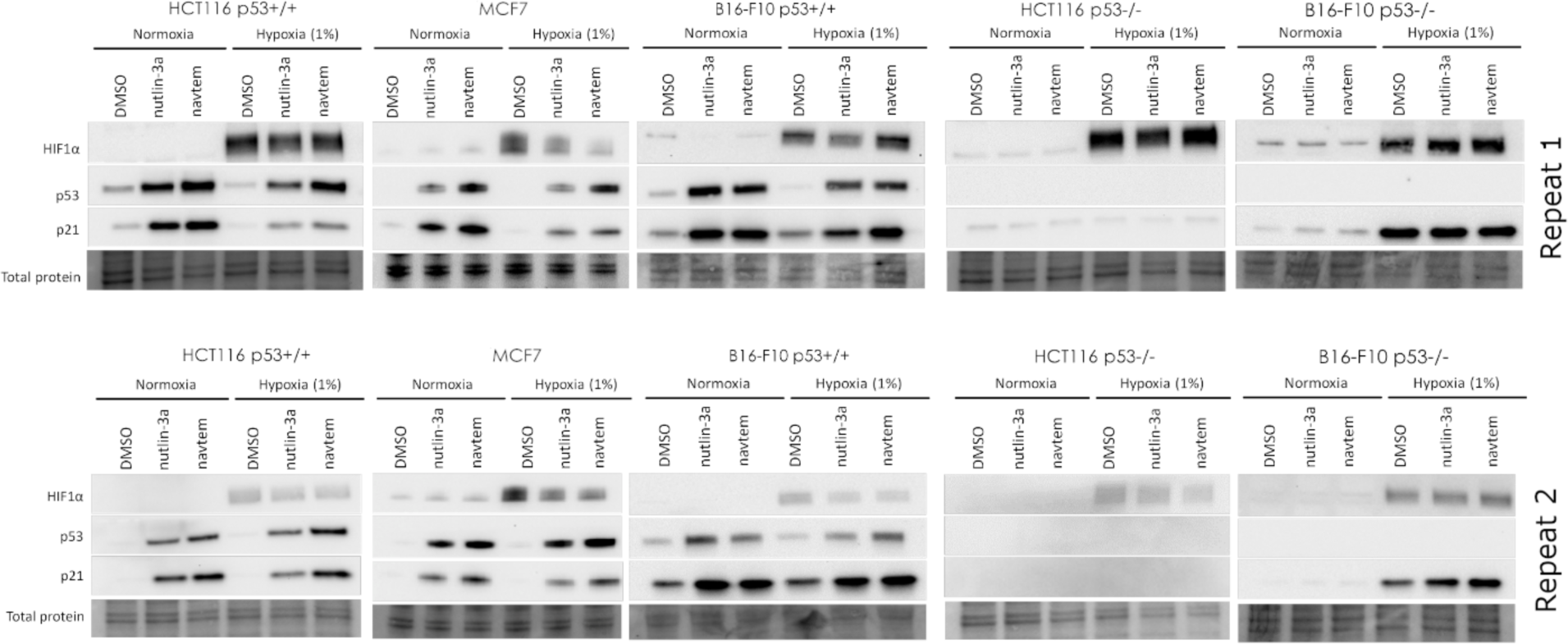
At the protein level, MDM2 inhibitors activate the p53-p21 pathway in hypoxic p53^wt^ cancer cells. The levels of HIF-1α, p53 and p21 levels were visualized after 24 h treatment with DMSO, nutlin-3a and navtemadlin (abbreviated navtem in figure) in either normoxia or hypoxia. Blots are from two independent experiments. Total protein images are shown as loading control.

### 3.3. Nutlin-3a and navtemadlin reduce the growth of p53^wt^ spheroids comprising innate hypoxia

Many studies have shown that 3D cancer models mimic many features of solid tumors, including variations in oxygen levels [30,31], gene expression [32,33], metabolism [34], and drug response profiles [32,33,35,36]. We therefore tested whether nutlin-3a and navtemadlin could reduce cell growth in tumor spheroids, a 3D *in vitro* model that naturally develops regions of hypoxia and necrosis at spheroid diameters of 400-600 μm due to an oxygen diffusion gradient [30,37,38]. To this end, we generated tumor spheroids from the p53^wt^ cell lines and treated them with the inhibitors when spheroids reached a diameter (> 500-550 μm). Both nutlin-3a and navtemadlin suppressed the growth of p53^WT^ spheroids from human cell lines (HCT116 p53^+/+^ and MCF7) by >75% when compared to DMSO. However, neither drug significantly reduced the growth of mouse p53^WT^ spheroids (Fig 5a-b). We observed that inhibitor treatment had varying effects on spheroid intactness depending on the cell line. While nearly all HCT116 p53^+/+^ spheroids shrunk from their starting size after treatment, a solid mass of ~450-500 μm in diameter remained (Fig 5a). In contrast, MCF7 spheroids were primarily cell debris (Fig 5a). B16-F10 p53^+/+^ spheroids appeared to be intact by brightfield microscopy but broke apart easily into cell debris upon pipetting. Upon close-up visualization of these B16-F10 spheroids, we noticed blebbing on the spheroid surface suggesting that cells may have been undergoing apoptosis (Fig 5c). As expected, no growth suppression was observed with inhibitor treatment in p53^KO^ spheroids (Fig 5d-e). Treatment with staurosporine, a non p53-specific inducer of apoptosis, prevented both p53^WT^ and p53^KO^ spheroids from growing (Fig 5d-e). These findings confirm that MDM2 inhibitors significantly reduce the growth of human, but not mouse, p53^WT^ cancer spheroids comprising innate hypoxia.

**Figure 5.**
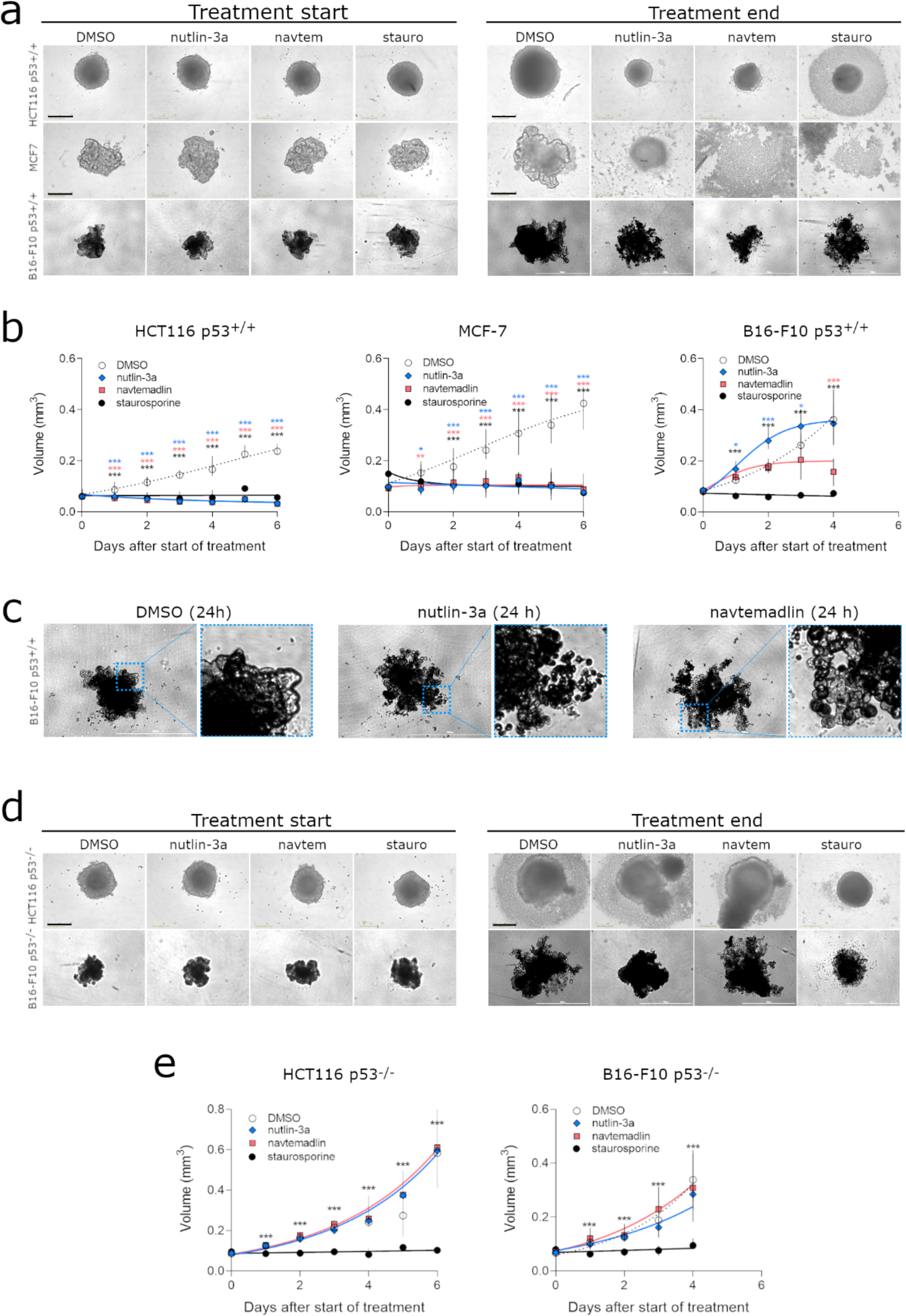
MDM2 inhibitors reduce growth of human p53^wt^ cells cultured as spheroids. (**a**) Brightfield images of HCT116 p53^+/+^, MCF7 and B16-F10 p53^+/+^ spheroids at start and end of treatment. (**b**) Growth curves of p53^wt^ spheroids treated with DMSO, nutlin-3a (human cells: 5 μM; mouse cells: 10 μM), navtemadlin (human cells: 1 μM; mouse cells: 2 μM), and staurosporine (human cells: 2 μM; mouse cells: 5 μM). (**c**) Brightfield images of B16-F10 p53^+/+^ spheroids after 24 h treatment with DMSO, nutlin-3a and navtemadlin. (**d**) Brightfield images of treated HCT116 p53^−/−^ and B16F10 p53^−/−^ spheroids at start and end of treatment. (**e**) Growth curves of HCT116 p53^−/−^ and B16F10 p53^−/−^ spheroids treated with DMSO, nutlin-3a, navtemadlin and staurosporine. Each data point represents a single spheroid (n = 6-18 spheroids/treatment) from 2-3 independent experiments. Statistical significance was assessed by mixed-effects ANOVA with matching followed by post-hoc Dunnett’s correction for multiple testing (unpaired, two-tailed, α = 0.05). Adjusted *P*-values from the Dunnett’s test are indicated (\**P* < 0.05, \*\**P* < 0.01, \*\*\**P* < 0.001).

## 4. Discussion

Our results show that the MDM2 inhibitors, nutlin-3a and navtemadlin, induce wild-type p53 in cancer cell lines cultured in hypoxia. Contrary to what we expected, the inhibitors reduced the growth of human and mouse p53wt cells in hypoxia through activation of the p53-p21 axis. The concentrations (IC_50_ values) required to induce these effects were not significantly different between normoxic and hypoxic conditions, and were consistent with values reported in previous studies performed in normoxia [6,39]. We also found that the inhibitors induced growth arrest in human cancer cells grown as 3D spheroid models comprising innate hypoxia, suggesting that the inhibitors retain efficacy even when cells are hypoxic prior to treatment. As expected, amongst the two inhibitors, navtemadlin was more potent than nutlin-3a at inducing a p53 response in all p53^WT^ cells cultured in hypoxia. Taken together, these findings indicate that the efficacy and potency of MDM2 inhibitors are not reduced in p53^WT^ cells cultured in hypoxia.

Although the potency and efficacy of MDM2 inhibitors were not altered in hypoxia, the efficacy varied by species, as evidenced in three ways. First, lower concentrations of inhibitors were required to elicit a p53 response in human cells than in mouse cells used in this study. Second, treatment with the inhibitors induced transcriptional activity by more than 9-fold in human cells, but by only 3-fold in mouse cells. Third, treatment with the inhibitors significantly reduced the volume of spheroids comprising human cells, but not that of spheroids comprising mouse cells; however, the inhibitors did affect the integrity and morphology of the spheroids comprising mouse cells. Although we do not know the exact reason for this variation across species, we speculate that differences in expression of p14ARF (or p19Arf in mice) among the cell lines may affect the efficacy of MDM2 inhibitors. Previous studies show that p19Arf can prevent p53 degradation through inactivation of MDM2’s ligase activity [40,41]. Given that the mouse B16-F10 cell line is p19Arf deficient [42], higher concentrations of inhibitors may be required to “bind” all the free MDM2 and stabilize p53. This explanation is consistent with previous reports showing that cell lines deficient in p19Arf are less able to elicit a p53 response [42].

Hypoxia alone also had varying effects by species. In the human p53^WT^ and p53^KO^ cancer cells studied here, 24 h exposure to hypoxia (1% O_2_) itself did not measurably induce cell cycle arrest, as evidenced by the lack of p21 expression in Western blots and by the lack of differences in S phase cells in hypoxia. By contrast, the same settings of hypoxia induced cell cycle arrest in the mouse p53^WT^ and p53^KO^ cell lines, as evidenced by the drop in S phase fraction of hypoxic cells. These phenotypic differences are consistent with those reported in the literature—some cell lines arrest in hypoxia, while others do not [22,43,44]. One potential mechanistic explanation is that p14ARF/p19Arf expression may protect cells from hypoxia-induced arrest, as p14ARF has been shown to inhibit HIF1α transcriptional activity [45]. The phenotypic differences may also arise from other variables such as the length of hypoxic exposure (ranging from 24 h to 72 h), oxygen levels (ranging from 0.02% to 1.4%), and cell seeding densities that influence gene expression [22,46].

Could MDM2 inhibitors induce p53 response in hypoxic regions of solid tumors? We found that while the inhibitors prevented the growth of 3D tumor spheroids, they did not eliminate them for some cell lines (e.g., HCT116). We hypothesize that this result could be due to cell cycle arrest or quiescence induced by hypoxia within the 3D spheroid [31,33,37], as long periods of hypoxia have been shown to arrest cells or to induce quiescence. Solid tumors also have acute regions of hypoxia (resulting from limited blood flow) and chronic regions of hypoxia (resulting from limited oxygen perfusion) [14]. Given that p53 is transcriptionally active only in proliferating cells [47], we expect therefore that MDM2 inhibitors would have reduced efficacy in arrested or quiescent cells. As such, we reason that the inhibitors would likely elicit a response in cancer cells experiencing acute hypoxia, but not in those experiencing chronic hypoxia. Further studies are required to dissect responses in chronic hypoxia.

## Acknowledgements

This research was supported by the Swedish Research Council awarded to DPL (grant number 2013-08807) and the Gunvor and Josef Anérs Foundation awarded to PK (grant number FB20-0092). BV was supported by the European Regional Development Fund - Project ENOCH (No. CZ.02.1.01/0.0/0.0/16_019/0000868). FW was supported by the Swedish Cancer Society and the Swedish Research Council. We thank the Biomedicum Flow Core Facility (Karolinska Institutet) for support and assistance with flow cytometers.

## Author contributions

Conception and design: DPL, PK

Data acquisition: ALC, PLB, KI, CB, SKS, PK

Analysis and interpretation of data: ALC, PLB, PK

Provided reagents and cell lines: LJ, FW, BV

Writing and revision of initial manuscript: ALC, PLB, DPL, PK

Revision of manuscript: ALC, PLB, KI, CB, SKS, LJ, FW, BV, DPL, PK

Study supervision and funding: DPL, PK

## Data availability

The datasets generated during the current study are available in the Zenodo repository (DOI: web link to be made available). Any other relevant data are available upon reasonable request from the corresponding authors.

## References

[1] A.M. Boutelle, L.D. Attardi, p53 and Tumor Suppression: It Takes a Network, Trends Cell Biol. 31 (2021) 298–310. https://doi.org/10.1016/j.tcb.2020.12.011.

[2] L. Bouaoun, D. Sonkin, M. Ardin, M. Hollstein, G. Byrnes, J. Zavadil, M. Olivier, TP53 Variations in Human Cancers: New Lessons from the IARC TP53 Database and Genomics Data, Hum. Mutat. 37 (2016) 865–876. https://doi.org/10.1002/humu.23035.

[3] C.J. Brown, S. Lain, C.S. Verma, A.R. Fersht, D.P. Lane, Awakening guardian angels: Drugging the P53 pathway, Nat. Rev. Cancer. 9 (2009) 862–873. https://doi.org/10.1038/nrc2763.

[4] A. Burgess, K.M. Chia, S. Haupt, D. Thomas, Y. Haupt, E. Lim, Clinical Overview of MDM2/X-Targeted Therapies, Front. Oncol. 6 (2016) 1–7. https://doi.org/10.3389/fonc.2016.00007.

[5] D. Spiegelberg, A.C. Mortensen, S. Lundsten, C.J. Brown, D.P. Lane, M. Nestor, The MDM2/MDMX-p53 antagonist PM2 radiosensitizes wild-type p53 tumors, Cancer Res. 78 (2018) 5084–93. https://doi.org/10.1158/0008-5472.CAN-18-0440.

[6] L. Vassilev, B.T. Vu, B. Graves, D. Carvajal, F. Podlaski, Z. Filipovic, N. Kong, U. Kammlott, C. Lukacs, C. Klein, N. Fotouhi, E.A. Liu, In Vivo Activation of the p53 Pathway by Small-Molecule Antagonists of MDM2, Science (80-.). 303 (2004) 844–849.

[7] L.R. Werner, S. Huang, D.M. Francis, E.A. Armstrong, F. Ma, C. Li, G. Iyer, J. Canon, P.M. Harari, Small molecule inhibition of MDM2-p53 interaction augments radiation response in human tumors, Mol. Cancer Ther. 14 (2015) 1994–2003. https://doi.org/10.1158/1535-7163.MCT-14-1056-T.

[8] A. Lakoma, E. Barbieri, S. Agarwal, J. Jackson, Z. Chen, Y. Kim, M. McVay, J.M. Shohet, E.S. Kim, The MDM2 small-molecule inhibitor RG7388 leads to potent tumor inhibition in p53 wild-type neuroblastoma, Cell Death Discov. 1 (2015) 1–9. https://doi.org/10.1038/cddiscovery.2015.26.

[9] N.G. Her, J.W. Oh, Y.J. Oh, S. Han, H.J. Cho, Y. Lee, G.H. Ryu, D.H. Nam, Potent effect of the MDM2 inhibitor AMG232 on suppression of glioblastoma stem cells, Cell Death Dis. 9 (2018) 1–12. https://doi.org/10.1038/s41419-018-0825-1.

[10] E. Barbieri, P. Mehta, Z. Chen, L. Zhang, A. Slack, S. Berg, J.M. Shohet, MDM2 inhibition sensitizes neuroblastoma to chemotherapy-induced apoptotic cell death, Mol. Cancer Ther. 5 (2006) 2358–2365. https://doi.org/10.1158/1535-7163.MCT-06-0305.

[11] D. Sun, Z. Li, Y. Rew, M. Gribble, M.D. Bartberger, H.P. Beck, J. Canon, A. Chen, X. Chen, D. Chow, J. Duquette, J. Eksterowicz, B. Fisher, B.M. Fox, J. Fu, A.Z. Gonzalez, F.G. De Turiso, J.B. Houze, X. Huang, M. Jiang, L. Jin, F. Kayser, J.J. Liu, M. Lo, A.M. Long, B. Lucas, L.R. Mcgee, J. Mcintosh, J. Mihalic, J.D. Oliner, T. Osgood, M.L. Peterson, P. Roveto, A.Y. Saiki, P. Sha, M. Toteva, Y. Wang, Y.C. Wang, S. Wortman, P. Yakowec, X. Yan, Q. Ye, D. Yu, M. Yu, X. Zhao, J. Zhou, J. Zhu, S.H. Olson, J.C. Medina, Discovery of AMG 232, a Potent, Selective, and Orally Bioavailable MDM2 - p53 Inhibitor in Clinical Development, J Med Chem. 57 (2014) 1454–1472. https://doi.org/10.1021/jm401753e.

[12] M.N. Nguyen, N. Sen, M. Lin, T.L. Joseph, C. Vaz, V. Tanavde, L. Way, T. Hupp, C.S. Verma, M.S. Madhusudhan, Discovering Putative Protein Targets of Small Molecules: A Study of the p53 Activator Nutlin, J. Chem. Inf. Model. 59 (2019) 1529–1546. https://doi.org/10.1021/acs.jcim.8b00762.

[13] L. Haronikova, O. Bonczek, P. Zatloukalova, F. Kokas-Zavadil, M. Kucerikova, P.J. Coates, R. Fahraeus, B. Vojtesek, Resistance mechanisms to inhibitors of p53-MDM2 interactions in cancer therapy: can we overcome them?, BioMed Central, 2021. https://doi.org/10.1186/s11658-021-00293-6.

[14] J.M. Brown, W.R. Wilson, Exploiting tumour hypoxia in cancer treatment, Nat. Rev. Cancer. 4 (2004) 437–447. https://doi.org/10.1038/nrc1367.

[15] J.P. Cosse, M. Ronvaux, N. Ninane, M.J. Raes, C. Michiels, Hypoxia-induced decrease in p53 protein level and increase in c-jun DNA binding activity results in 18 cancer cell resistance to etoposide, Neoplasia. 11 (2009) 976–986. https://doi.org/10.1593/neo.09632.

[16] S. Strese, M. Fryknäs, R. Larsson, J. Gullbo, Effects of hypoxia on human cancer cell line chemosensitivity, BMC Cancer. 13 (2013). https://doi.org/10.1186/1471-2407-13-331.

[17] A.L. Harris, Hypoxia - A key regulatory factor in tumour growth, Nat. Rev. Cancer. 2 (2002) 38–47. https://doi.org/10.1038/nrc704.

[18] A. Nijhuis, H. Thompson, J. Adam, A. Parker, L. Gammon, A. Lewis, J.G. Bundy, T. Soga, A. Jalaly, D. Propper, R. Jeffery, N. Suraweera, S. McDonald, M.A. Thaha, R. Feakins, R. Lowe, C.L. Bishop, A. Silver, Remodelling of microRNAs in colorectal cancer by hypoxia alters metabolism profiles and 5-fluorouracil resistance, Hum. Mol. Genet. 26 (2017) 1552–1564. https://doi.org/10.1093/hmg/ddx059.

[19] Z. Jiang, J. Yang, A. Dai, Y. Wang, W. Li, Z. Xie, Ribosome profiling reveals translational regulation of mammalian cells in response to hypoxic stress, BMC Genomics. 18 (2017) 1–12. https://doi.org/10.1186/s12864-017-3996-8.

[20] V. Marcel, F. Catez, J.J. Diaz, P53, a translational regulator: Contribution to its tumour-suppressor activity, Oncogene. 34 (2015) 5513–5523. https://doi.org/10.1038/onc.2015.25.

[21] T.G. Graeber, C. Osmanian, T. Jackstt, D.E. Housmant, C.J. Koch, S.W. Lowetll, A.J.G. Ii, Hypoxia-mediated selection of cells with diminished apoptotic potential in Solid Tumours, Nature. 379 (1996) 88–91.

[22] B. Ortmann, J. Druker, S. Rocha, Cell cycle progression in response to oxygen levels, Cell. Mol. Life Sci. 71 (2014) 3569–3582. https://doi.org/10.1007/s00018-014-1645-9.

[23] V. Bhandari, C. Hoey, L.Y. Liu, E. Lalonde, J. Ray, J. Livingstone, R. Lesurf, Y.J. Shiah, T. Vujcic, X. Huang, S.M.G. Espiritu, L.E. Heisler, F. Yousif, V. Huang, T.N. Yamaguchi, C.Q. Yao, V.Y. Sabelnykova, M. Fraser, M.L.K. Chua, T. van der Kwast, S.K. Liu, P.C. Boutros, R.G. Bristow, Molecular landmarks of tumor hypoxia across cancer types, Nat. Genet. 51 (2019) 308–318. https://doi.org/10.1038/s41588-018-0318-2.

[24] J. Yang, A. Ahmed, E. Poon, N. Perusinghe, A. de Haven Brandon, G. Box, M. Valenti, S. Eccles, K. Rouschop, B. Wouters, M. Ashcroft, Small-Molecule Activation of p53 Blocks Hypoxia-Inducible Factor 1 and Vascular Endothelial Growth Factor Expression In Vivo and Leads to Tumor Cell Apoptosis in Normoxia and Hypoxia, Mol. Cell. Biol. 29 (2009) 2243–2253. https://doi.org/10.1128/MCB.00959-08.

[25] A. Weilbacher, M. Gutekunst, M. Oren, W.E. Aulitzky, H. Van Der Kuip, RITA can induce cell death in p53-defective cells independently of p53 function via activation of JNK/SAPK and p38, Cell Death Dis. 5 (2014) e1318-11. https://doi.org/10.1038/cddis.2014.284.

[26] J. De Lange, L. V. Ly, K. Lodder, M. Verlaan-De Vries, A.F.A.S. Teunisse, M.J. Jager, A.G. Jochemsen, Synergistic growth inhibition based on small-molecule p53 activation as treatment for intraocular melanoma, Oncogene. 31 (2012) 1105–1116. https://doi.org/10.1038/onc.2011.309.

[27] K. Ingelshed, D. Spiegelberg, P. Kannan, L. Pavenius, L. Jiang, S. Eisinger, D. Lianoudaki, J. Hacheney, D. Lama, C. Bosdotter, N. Fritz, M.C.I. Karlsson, F. Wermeling, M. Nestor, F. Castillo, W.W. Kretzschmar, O. Al-Radi, E. Villablanca, D.P. Lane, S. Sedimbi, The MDM2 inhibitor AMG 232 arrests mouse tumor growth in vivo and potentiates radiotherapy, Press. Cancer Res Commun. (2022).

[28] K. Brimacombe, M. Hall, D. Auld, J. Inglese, C. Austin, M. Gottesman, K. Fung, A dual-fluorescence high-throughput cell line system for probing multidrug resistance, Assay Drug Dev. Tech. 7 (2009) 233–49.

[29] W. Chen, C. Wong, E. Vosburgh, A.J. Levine, D.J. Foran, E.Y. Xu, B. Sackler, E. High, High-throughput Image Analysis of Tumor Spheroids: A User-friendly Software Application to Measure the Size of Spheroids Automatically and Accurately, J. Vis. Exp. 89 (2014) 51639. https://doi.org/10.3791/51639.

[30] D.R. Grimes, P. Kannan, A. McIntyre, A. Kavanagh, A. Siddiky, S. Wigfield, A. Harris, M. Partridge, The role of oxygen in avascular tumor growth, PLoS One. 11 (2016) 1–19. https://doi.org/10.1371/journal.pone.0153692.

[31] F. Hirschhaeuser, H. Menne, C. Dittfeld, J. West, W. Mueller-Klieser, L. a. Kunz-Schughart, Multicellular tumor spheroids: An underestimated tool is catching up again, J. Biotechnol. 148 (2010) 3–15. https://doi.org/10.1016/j.jbiotec.2010.01.012.

[32] V. Härmä, J. Virtanen, R. Mäkelä, A. Happonen, J.-P. Mpindi, M. Knuuttila, P. Kohonen, J. Lötjönen, O. Kallioniemi, M. Nees, A comprehensive panel of threedimensional models for studies of prostate cancer growth, invasion and drug responses., PLoS One. 5 (2010) e10431. https://doi.org/10.1371/journal.pone.0010431.

[33] A.C. Luca, S. Mersch, R. Deenen, S. Schmidt, I. Messner, K.L. Schäfer, S.E. Baldus, W. Huckenbeck, R.P. Piekorz, W.T. Knoefel, A. Krieg, N.H. Stoecklein, Impact of the 3D Microenvironment on Phenotype, Gene Expression, and EGFR Inhibition of Colorectal Cancer Cell Lines, PLoS One. 8 (2013). https://doi.org/10.1371/journal.pone.0059689.

[34] S. Russell, J. Wojtkowiak, A. Neilson, R.J. Gillies, Metabolic Profiling of healthy and cancerous tissues in 2D and 3D, Sci. Rep. 7 (2017) 1–11. https://doi.org/10.1038/s41598-017-15325-5.

[35] A. Riedl, M. Schlederer, K. Pudelko, M. Stadler, S. Walter, D. Unterleuthner, C. Unger, N. Kramer, M. Hengstschläger, L. Kenner, D. Pfeiffer, G. Krupitza, H. Dolznig, Comparison of cancer cells in 2D vs 3D culture reveals differences in AKT-mTOR-S6K signaling and drug responses, J. Cell Sci. 130 (2017) 203–218. https://doi.org/10.1242/jcs.188102.

[36] Y. Imamura, T. Mukohara, Y. Shimono, Y. Funakoshi, N. Chayahara, M. Toyoda, N. Kiyota, S. Takao, S. Kono, T. Nakatsura, H. Minami, Comparison of 2D- and 3D-culture models as drug-testing platforms in breast cancer, Oncol. Rep. 33 (2015) 1837–1843. https://doi.org/10.3892/or.2015.3767.

[37] S. Riffle, R.N. Pandey, M. Albert, R.S. Hegde, Linking hypoxia, DNA damage and proliferation in multicellular tumor spheroids, BMC Cancer. 17 (2017) 1–12. https://doi.org/10.1186/s12885-017-3319-0.

[38] D.R. Grimes, C. Kelly, K. Bloch, M. Partridge, A method for estimating the oxygen consumption rate in multicellular tumour spheroids, J. R. Soc. Interface. 11 (2014). https://doi.org/10.1098/rsif.2013.1124.

[39] J. Canon, T. Osgood, S.H. Olson, A.Y. Saiki, R. Robertson, D. Yu, J. Eksterowicz, Q. Ye, L. Jin, A. Chen, J. Zhou, D. Cordover, S. Kaufman, R. Kendall, J.D. Oliner, A. Coxon, R. Radinsky, The MDM2 inhibitor AMG 232 demonstrates robust antitumor efficacy and potentiates the activity of p53-inducing cytotoxic agents, Mol. Cancer Ther. 14 (2015) 649–658. https://doi.org/10.1158/1535-7163.MCT-14-0710.

[40] R. Honda, H. Yasuda, Association of p19(ARF) with Mdm2 inhibits ubiquitin ligase activity of Mdm2 for tumor suppressor p53, EMBO J. 18 (1999) 22–27. https://doi.org/10.1093/emboj/18.1.22.

[41] J. Pomerantz, N. Schreiber-Agus, N.J. Liégeois, A. Silverman, L. Alland, L. Chin, J. Potes, K. Chen, I. Orlow, H.W. Lee, C. Cordon-Cardo, R.A. DePinho, The Ink4a tumor suppressor gene product, p19(Arf), interacts with MDM2 and neutralizes MDM2’s inhibition of p53, Cell. 92 (1998) 713–723. https://doi.org/10.1016/S0092-8674(00)81400-2.

[42] C.A. Merkel, R.B. da Silva Soares, A.C. V. de Carvalho, D.B. Zanatta, M.C. Bajgelman, P. Fratini, E. Costanzi-Strauss, B.E. Strauss, Activation of endogenous p53 by combined p19Arf gene transfer and nutlin-3 drug treatment modalities in the murine cell lines B16 and C6, BMC Cancer. 10 (2010). https://doi.org/10.1186/1471-2407-10-316.

[43] A.H. Box, D.J. Demetrick, Cell cycle kinase inhibitor expression and hypoxia-induced cell cycle arrest in human cancer cell lines, Carcinogenesis. 25 (2004) 2325–2335. https://doi.org/10.1093/carcin/bgh274.

[44] S. Yoshiba, D. Ito, T. Nagumo, T. Shirota, M. Hatori, S. Shintani, Hypoxia induces 22 resistance to 5-fluorouracil in oral cancer cells via G1 phase cell cycle arrest, Oral Oncol. 45 (2009) 109–115. https://doi.org/10.1016/j.oraloncology.2008.04.002.

[45] K. Fatyol, A.A. Szalay, The p14ARF Tumor Suppressor Protein Facilitates Nucleolar Sequestration of Hypoxia-inducible Factor-1α (HIF-1α) and Inhibits HIF-1-mediated Transcription, J. Biol. Chem. 276 (2001) 28421–28429. https://doi.org/10.1074/jbc.M102847200.

[46] L. LeBlanc, M. Kim, A. Kambhampati, A.J. Son, N. Ramirez, J. Kim, Β-Catenin Links Cell Seeding Density To Global Gene Expression During Mouse Embryonic Stem Cell Differentiation, IScience. 25 (2022) 103541. https://doi.org/10.1016/j.isci.2021.103541.

[47] Y. Xue, N. Barker, S. Hoon, P. He, T. Thakur, S.R. Abdeen, P. Maruthappan, F.J. Ghadessy, D.P. Lane, Bortezomib stabilizes and activates p53 in proliferative compartments of both normal and tumor tissues in vivo, Cancer Res. 79 (2019) 3595–3607. https://doi.org/10.1158/0008-5472.CAN-18-3744.

[48] B. Vojtěšek, J. Bártek, C.A. Midgley, D.P. Lane, An immunochemical analysis of the human nuclear phosphoprotein p53. New monoclonal antibodies and epitope mapping using recombinant p53, J. Immunol. Methods. 151 (1992) 237–244. https://doi.org/10.1016/0022-1759(92)90122-A.

